## supplementary Figures S1-S11 for "Control of innate olfactory valence by segregated cortical amygdala circuits"

#### Figure S1

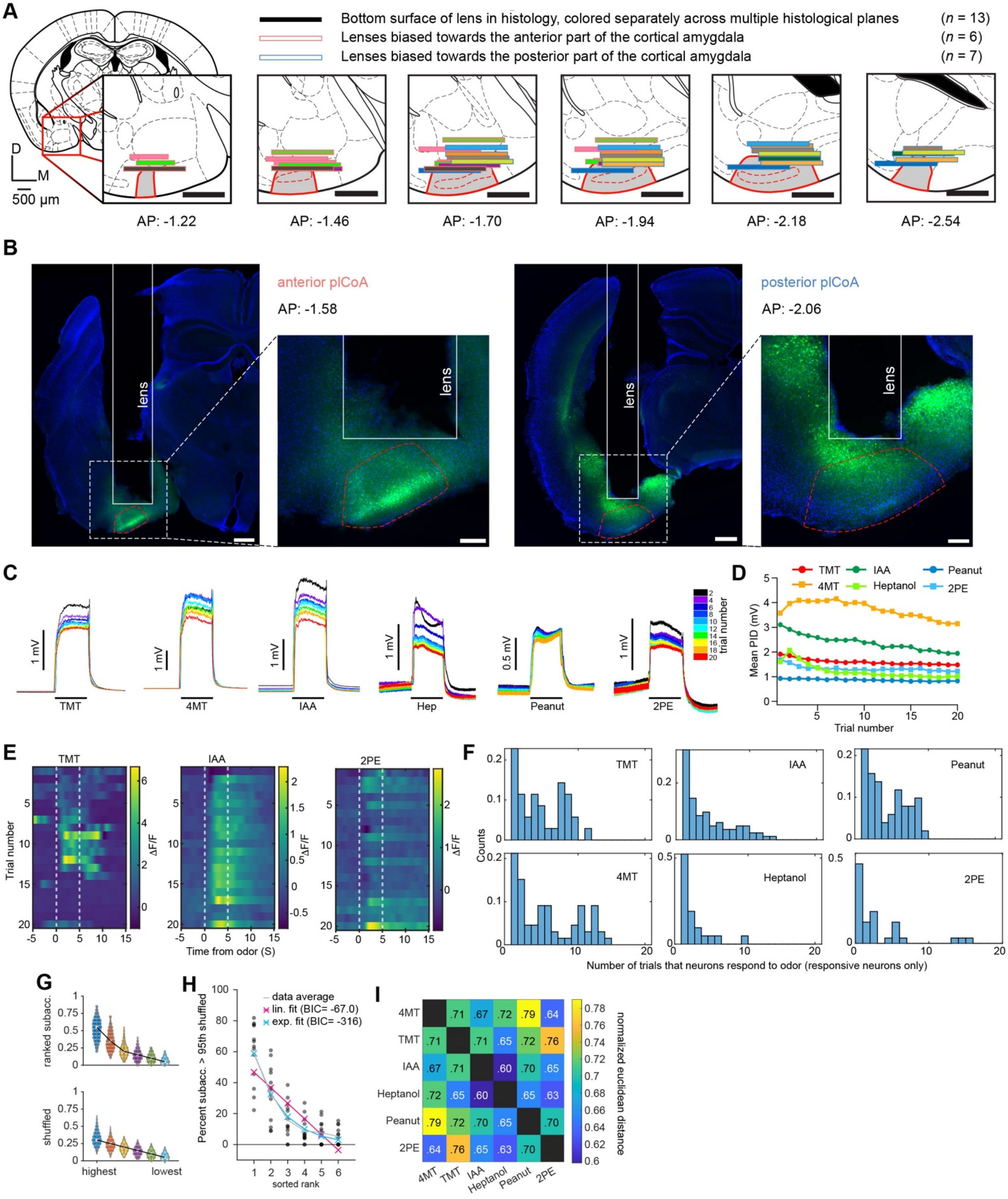
A
Bottom surface of lens in histology, colored separately across multiple histological planes (n = 13)
Lenses biased towards the anterior part of the cortical amygdala (n = 6)
Lenses biased towards the posterior part of the cortical amygdala (n = 7)
D
M
500 µm
AP: -1.22
AP: -1.46
AP: -1.70
AP: -1.94
AP: -2.18
AP: -2.54
B
anterior plCoA
AP: -1.58
lens
posterior plCoA
AP: -2.06
C
TMT
4MT
IAA
Hep
Peanut
2PE
trial number
D
TMT
IAA
Peanut
4MT
Heptanol
2PE
Mean PID (mV)
Trial number
E
TMT
IAA
2PE
Trial number
ΔF/F
Time from odor (S)
F
Counts
Number of trials that neurons respond to odor (responsive neurons only)
G
ranked subacc.
shuffled
highest
lowest
H
data average
lin. fit (BIC= -67.0)
exp. fit (BIC= -316)
Percent subacc. > 95th shuffled
sorted rank
I
4MT
TMT
IAA
Heptanol
Peanut
2PE
normalized euclidean distance

**Figure S1. Additional information for imaging experiments.** Related to **Figure 1.**
(**A**) Histologically verified placements for GRIN lenses (bars) in wild type animals infected with AAV-hSyn-GCaMP8s for calcium imaging experiments. Horizontal lines indicate locations where lens placements were visible in coronal sections at various positions in the plCoA anterior-posterior axis. Different color bars represent placements in each of 13 animals. Those scored as anterior are outlined in red, and those in posterior are outlined in blue.
**(B)** Histology for anterior plCoA (left) and posterior plCoA (right) lens implant and jGCaMP8s (green) viral injection sites in representative animals. Scale bar for full size images, 500 μm; scale bar for magnified images, 200 μm.
(**C**) PID recordings across 20 repeated presentations of each odor. Colored traces represent individual trials
(every other trial shown for clarity). Horizontal black bars indicate 5-second odor deliveries.
(**D**) Mean PID amplitude during the odor presentation period plotted across trials for each odor. For each trace,
the baseline was shifted to approximately zero to compensate for drift in recordings over time.
(**E**) Representative peristimulus time heatmaps showing single-neuron responses across repeated odor presentations. Each panel displays activity from one neuron to one odor. The odor delivery window (0–5 s) is indicated by white dashed lines.
(**F**) Histograms reporting the distribution for the number of trials neurons respond to each odor. Neurons were classified as responsive on any trial with a $Z > 2$.
(**G**) Ranked sub-accuracies for single-neuron MNR classifiers (Figure 1L) in violin plots for those trained on real (top) or shuffled (bottom) data. Black lines connect the median across each rank.
(**H**) Scatter plot showing proportion of sub-accuracies exceeding the 95th percentile of the shuffled controls for each animal, ordered by sub-accuracy rank. Red and blue lines show the linear and exponential fit, respectively, and a gray dashed line connects data averages.
**(I)** Heatmap of the normalized average pairwise Euclidean distance between odor response vectors across biological replicates.

**Figure S2**

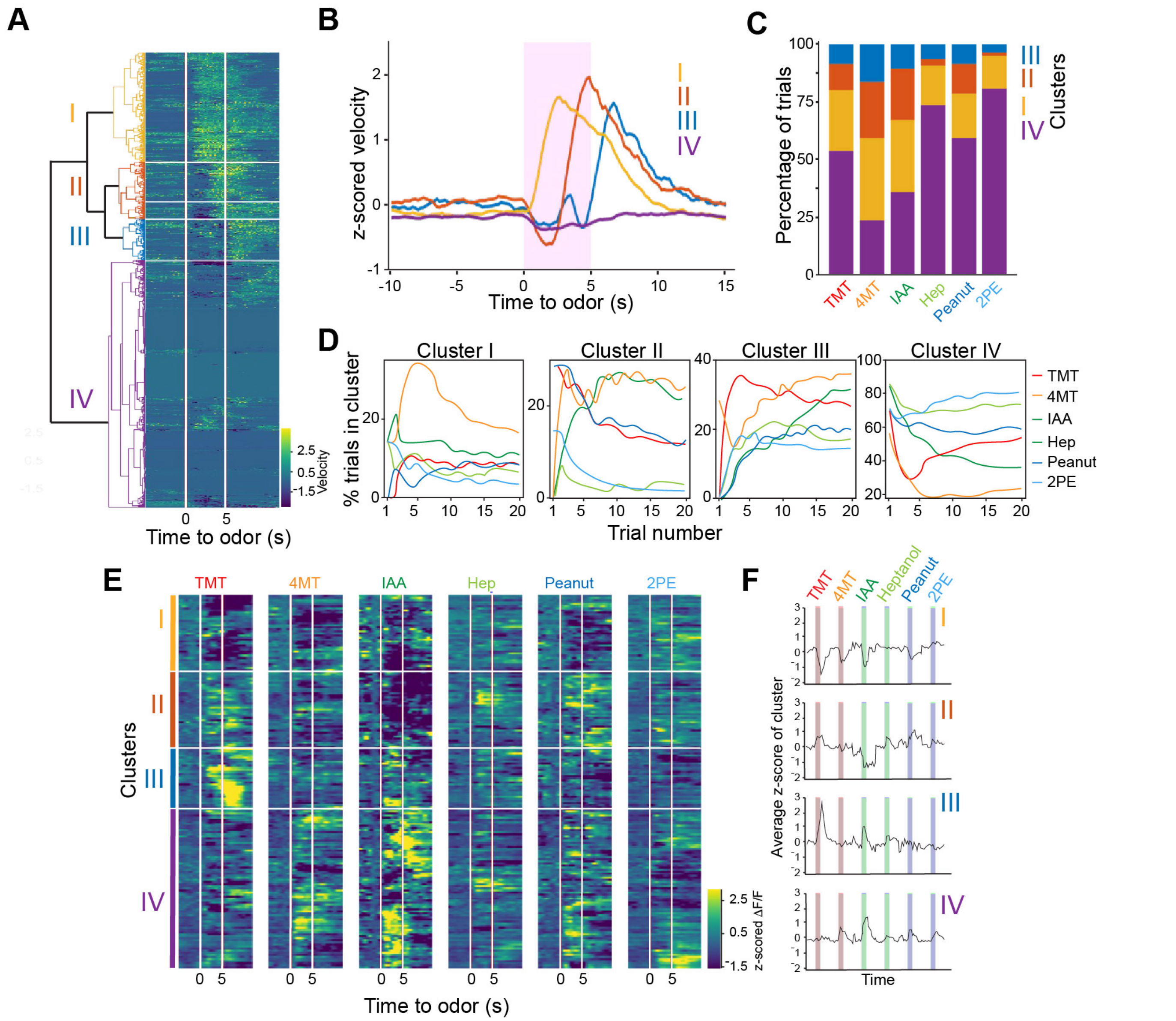


**Figure S2. Analysis of odor-evoked calcium activity with respect to walking behavior during imaging trials.** Related to **Figure 1.**

(**A**) Heatmap of normalized walking velocity in head-fixed mice across imaging trials. Trials grouped by hierarchical clustering of velocity during and after odor exposure, with the dendrogram (left). Odor delivery indicated by vertical red lines.

**(B)** Mean of trial-averaged and z-scored velocity for each cluster, without respect to odor identity.

(**C**) Percentage of trials for each cluster separated by odor.

(**D**) Percentage of trials for each cluster as a function of trial number, plotted for each odor.

(**E**) Heatmap of trial averaged and z-scored odor-evoked activity clustered according to the walking velocity cluster (A-D).

(**F**) Mean of trial-averaged and Z-scored calcium activity for each velocity cluster. concatenated. The order of color-coded blocks corresponds to the order of clusters in (E).

### Figure S3

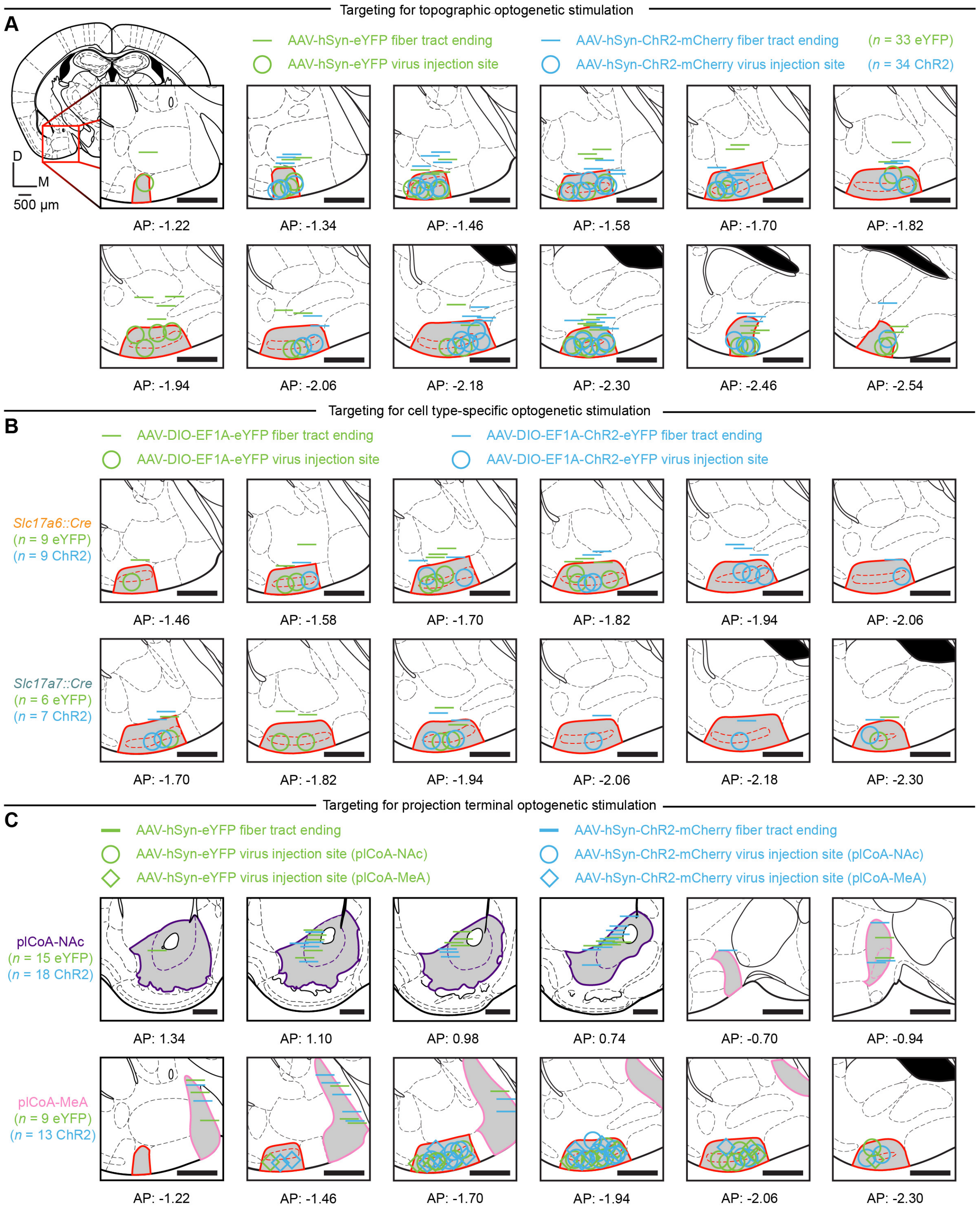
Targeting for topographic optogenetic stimulation
A
AAV-hSyn-eYFP fiber tract ending
AAV-hSyn-eYFP virus injection site
AAV-hSyn-ChR2-mCherry fiber tract ending
AAV-hSyn-ChR2-mCherry virus injection site
(n = 33 eYFP)
(n = 34 ChR2)
D
M
500 μm
AP: -1.22
AP: -1.34
AP: -1.46
AP: -1.58
AP: -1.70
AP: -1.82
AP: -1.94
AP: -2.06
AP: -2.18
AP: -2.30
AP: -2.46
AP: -2.54
Targeting for cell type-specific optogenetic stimulation
B
AAV-DIO-EF1A-eYFP fiber tract ending
AAV-DIO-EF1A-eYFP virus injection site
AAV-DIO-EF1A-ChR2-eYFP fiber tract ending
AAV-DIO-EF1A-ChR2-eYFP virus injection site
Slc17a6::Cre
(n = 9 eYFP)
(n = 9 ChR2)
AP: -1.46
AP: -1.58
AP: -1.70
AP: -1.82
AP: -1.94
AP: -2.06
Slc17a7::Cre
(n = 6 eYFP)
(n = 7 ChR2)
AP: -1.70
AP: -1.82
AP: -1.94
AP: -2.06
AP: -2.18
AP: -2.30
Targeting for projection terminal optogenetic stimulation
C
AAV-hSyn-eYFP fiber tract ending
AAV-hSyn-eYFP virus injection site (plCoA-NAc)
AAV-hSyn-eYFP virus injection site (plCoA-MeA)
AAV-hSyn-ChR2-mCherry fiber tract ending
AAV-hSyn-ChR2-mCherry virus injection site (plCoA-NAc)
AAV-hSyn-ChR2-mCherry virus injection site (plCoA-MeA)
plCoA-NAc
(n = 15 eYFP)
(n = 18 ChR2)
AP: 1.34
AP: 1.10
AP: 0.98
AP: 0.74
AP: -0.70
AP: -0.94
plCoA-MeA
(n = 9 eYFP)
(n = 13 ChR2)
AP: -1.22
AP: -1.46
AP: -1.70
AP: -1.94
AP: -2.06
AP: -2.30

**Figure S3. Targeting of plCoA neurons for optogenetic stimulation.** Related to **Figures 2**, **5**, **7**, **S3**, and **S8**.
(**A**) Histologically verified placements for optic fiber implants (bars) and viral injection sites (circles) in wild type animals infected with AAV-hSyn-ChR2-mCherry (blue) or AAV-hSyn-eYFP (green) in topographic optogenetic cell body stimulation experiments in **Figure 1** and **S2**.
(**B**) Same as (**A**), but for cell-type specific optogenetic cell body stimulation experiments for *Slc17a7::Cre* (top) and *Slc17a6::Cre* (bottom) animals infected with AAV-DIO-EF1A-ChR2-eYFP (blue) or AAV-EF1A-DIO-eYFP (green) in **Figure 5**.
(**C**) Respective placements for fiber implants (bars) and injection sites (plCoA-NAc, circles; plCoA-MeA, diamonds) in wild-type animals infected with AAV-hSyn-ChR2-mCherry (blue) or AAV-hSyn-eYFP (green) in projection-specific optogenetic axon terminal stimulation experiments in **Figure 7** and **S8.**
*n* denotes number of mice per group batched across 4-quad, elevated plus maze, and open field test experiments, exceeding *n* values for individual experiments due to behavioral cohort design (see STAR Methods). Relevant regions are highlighted in grey and outlined: plCoA (red), NAc (purple, only in **C**), and MeA (pink, only in **C**). All mouse brain sections reproduced from Paxinos and Franklin, 5$^{th}$ Edition, and numbers below all images denote its anterior-posterior distance from bregma in this atlas.[36] All scale bars, 500 µm.

**Figure S4**

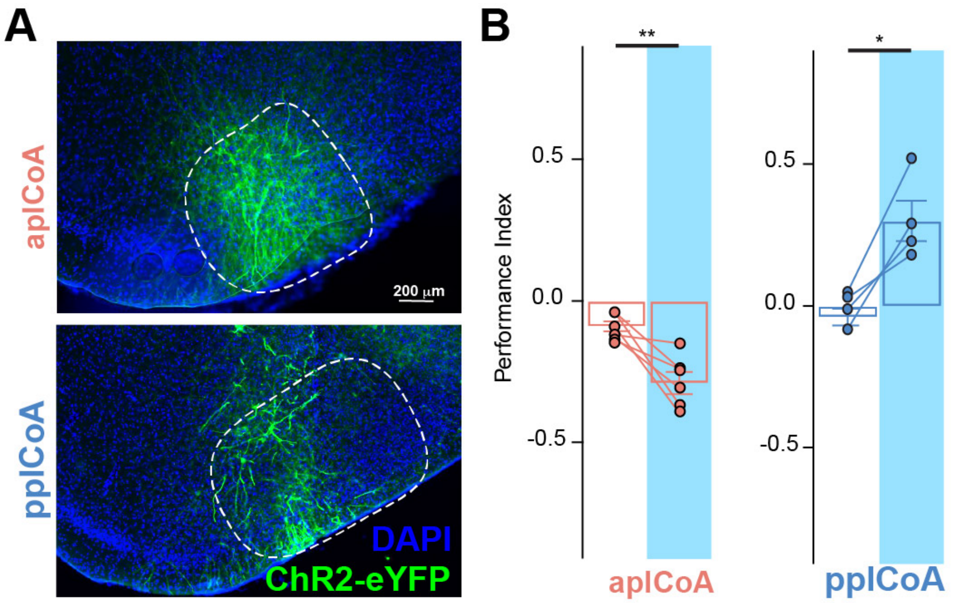


**Figure S4. Activation of sparse ensembles in apICoA and ppICoA elicits avoidance and approach**, Related to Figure 2.
The Arc-creER$^{T2}$ mouse [14] in combination with a cre-dependent AAV was used to express ChR2-eYFP in either apICoA or ppICoA. Mice were administered tamoxifen and exposed to TMT.
(A) Representative images of ChR2-eYFP in the apICoA (top) or ppICoA (bottom).
(B) Optogenetic stimulation-induced change in performance index for apICoA- (left) or ppICoA-injected (right) mice. Photostimulation induces aversive responses in apICoA, and approach responses in ppICoA. * $p< 0.05$; ** $p < 0.01$; Additional specific details of statistical tests can be found in Supplemental Table 1.

#### Figure S5

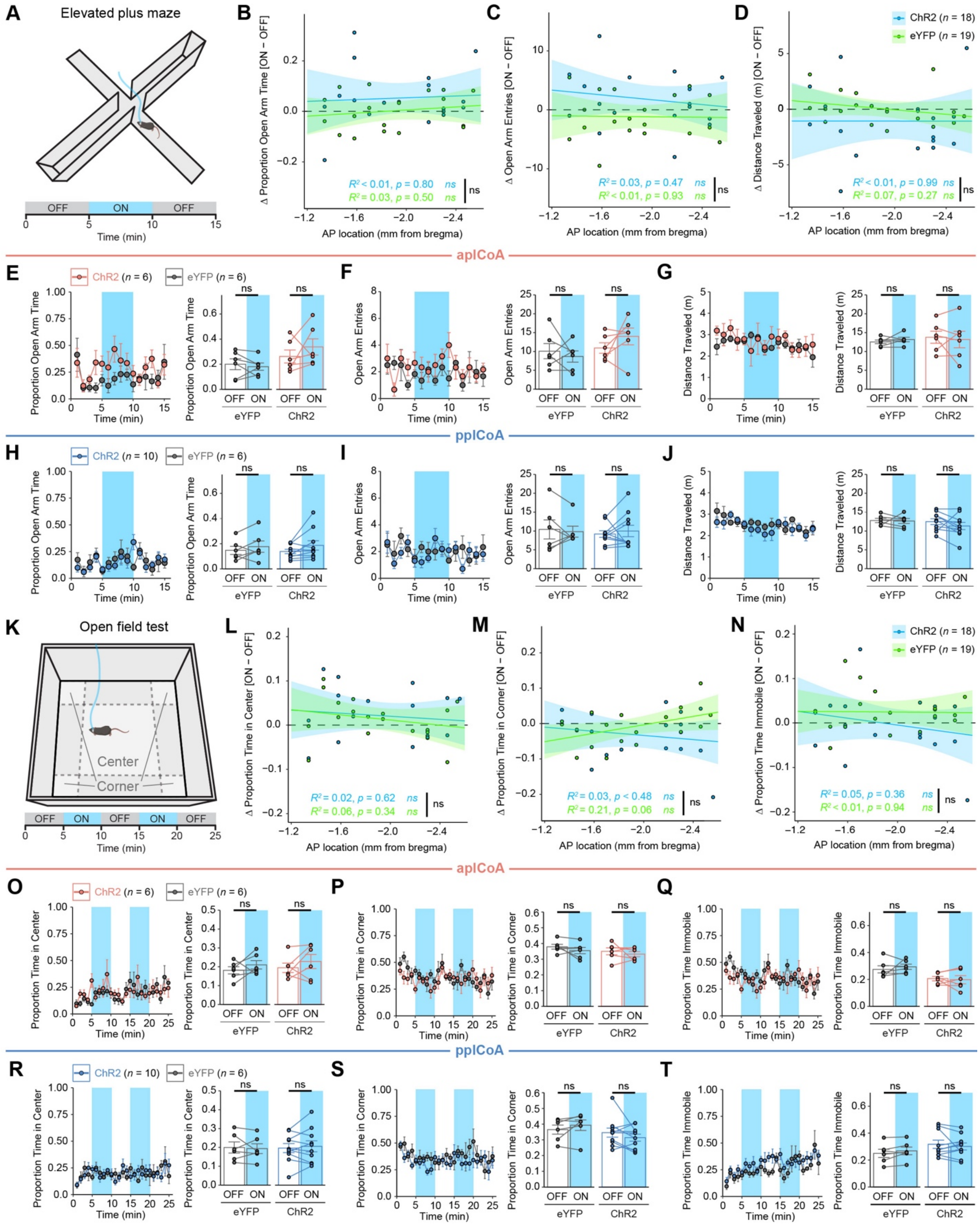
A
Elevated plus maze
OFF ON OFF
Time (min)
B
Δ Proportion Open Arm Time [ON – OFF]
AP location (mm from bregma)
$R^2$ < 0.01, p = 0.80 ns
$R^2$ = 0.03, p = 0.50 ns
ns
C
Δ Open Arm Entries [ON – OFF]
$R^2$ = 0.03, p = 0.47 ns
$R^2$ < 0.01, p = 0.93 ns
D
Δ Distance Traveled (m) [ON – OFF]
ChR2 (n = 18)
eYFP (n = 19)
$R^2$ < 0.01, p = 0.99 ns
$R^2$ = 0.07, p = 0.27 ns
apICoA
E
ChR2 (n = 6)
eYFP (n = 6)
Proportion Open Arm Time
OFF ON
eYFP ChR2
F
Open Arm Entries
G
Distance Traveled (m)
ppICoA
H
ChR2 (n = 10)
eYFP (n = 6)
I
J
K
Open field test
Center
Corner
OFF ON OFF ON OFF
L
Δ Proportion Time in Center [ON – OFF]
$R^2$ = 0.02, p = 0.62 ns
$R^2$ = 0.06, p = 0.34 ns
M
Δ Proportion Time in Corner [ON – OFF]
$R^2$ = 0.03, p < 0.48 ns
$R^2$ = 0.21, p = 0.06 ns
N
Δ Proportion Time Immobile [ON – OFF]
$R^2$ = 0.05, p = 0.36 ns
$R^2$ < 0.01, p = 0.94 ns
O
Proportion Time in Center
P
Proportion Time in Corner
Q
Proportion Time Immobile
R
S
T

**Figure S5. Behavioral effects of topographic plCoA stimulation are limited to valence alone.** Related to **Figure 2**.

**(A)** Behavioral paradigm for optogenetic stimulation in the open field test.

**(B-D)** Optogenetic stimulation-induced change in time spent (**B**) and number of entries (**C**) into the open arms, as well as distance traveled (**D**) in the elevated plus maze is not correlated to anteroposterior axis position in plCoA.

**(E-G)** Effects of optogenetic stimulation of aplCoA neurons in the elevated plus maze in 1 minute bins over time (left) and during off and on periods (right).Photostimulation of aplCoA neurons does not induce a significant change in time spent (**E**) and number of entries (**F**) into the open arms, as well as distance traveled (**G**) in the elevated plus maze.

**(H-J)** Effects of optogenetic stimulation of pplCoA neurons in the elevated plus maze in 1minute bins over time (left) and during off and on periods (right). Photostimulation of pplCoA neurons does not induce a significant change in time spent (**H**) and number of entries (**I**) into the open arms, as well as distance traveled (**J**) in the elevated plus maze.

**(K)** Behavioral paradigm for optogenetic stimulation in the open field test.

**(L-N)** Optogenetic stimulation-induced change in time spent in the center (**L**) and corners (**M**), as well as distance traveled (**N**) in the open field test is not correlated to anteroposterior axis position in plCoA.

**(O-Q)** Effects of optogenetic stimulation of aplCoA neurons in the open field test in 1 mintue time bins (left) and during off and on periods (right). Photostimulation of aplCoA neurons does not induce a significant change time spent in the center (**O**) and corners (**P**), as well as distance traveled (**Q**) in the open field test.

**(R-T)** Effects of optogenetic stimulation of pplCoA neurons in the open field test in 1 minute time bins (left) and during off and on periods (right). Photostimulation of pplCoA neurons does not induce a significant change time spent in the center (**R**) and corners (**S**), as well as distance traveled (**T**) in the open field test.

All “ON” and “OFF” comparisons in bar graphs and linear regressions are on a per 5-minute basis. **(B-D, L-M)** Least-squares linear regression ± 95% confidence interval. Across panels: ns, not significant. Additional specific details of statistical tests can be found in Supplemental Table 1.

#### Figure S6

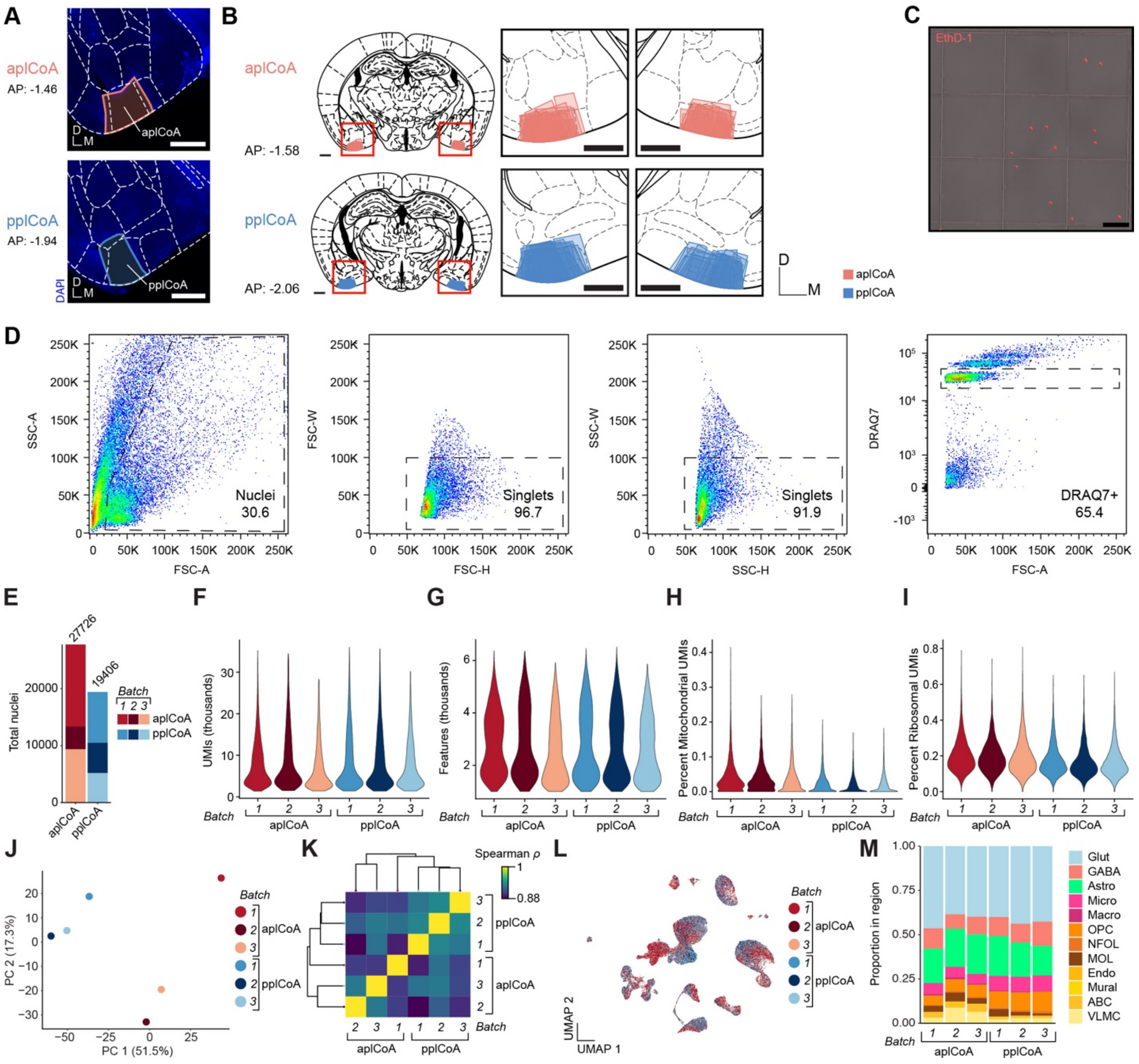


**Figure S6. Additional information and quality-control for single-nucleus sequencing experiments.** Related to **Figure 3**.

**(A)** Representative images of tissue microdissection sites from aplCoA and pplCoA following extraction and DAPI staining (blue). Scale bars, 500 µm.

**(B)** Location of all tissue sample sites for used for snRNA-seq, color coded by plCoA zone (n = 3 pools per zone, 4-11 sections per pool). Scale bars, 500 µm.

**(C)** Validation of nuclear enrichment after FANS. Ethidium homodimer-1 (EthD-1, red) labels nuclei on a hemocytometer after sorting, with an absence of non-nuclear, EthD-1-negative debris. Scale bar, 100 µm.

**(D)** Common gating strategy for FANS sorts for snRNA-seq in a representative sample.[102] Far left, morphology gate on forward and side scatter area excludes likely debris. Middle left,

forward scatter gate excludes nuclear doublets with high forward scatter width. Middle right, side scatter gate excludes nuclear multiplets with high side scatter width. Far right, stoichiometric DRAQ7+ fluorescence allows enrichment of single nuclei and exclusion of debris and multiplets.
**(E)** Absolute number and proportion of snRNA-seq nuclei passing quality control filters from each replicate in each plCoA zone (n = 27,726 in aplCoA, 19,406 in pplCoA, 3 libraries/batches each).
**(F)** Violin plot of UMIs detected per snRNA-seq nucleus for each replicate, filtered at the median per library + five times the median absolute deviation within each library (median 6081 UMIs/nucleus).
**(G)** Violin plot of genes detected per nucleus from each replicate, filtered at a minimum of 1000 features per nucleus (median 2547 genes/nucleus).
**(H)** Percent mitochondrial gene UMIs per snRNA-seq nucleus, filtered at median + five times the median absolute deviation per library (median 0.02% mitochondrial UMIs/nucleus).
**(I)** Percent ribosomal gene UMIs per snRNA-seq nucleus, filtered at median + five times the median absolute deviation per library (median 0.17% ribosomal UMIs/nucleus).
**(J)** Principal component analysis of pseudobulk snRNA-seq samples created from each batch, colored based on their combination of zone and batch identity.
**(K)** Evaluation of transcriptomic homology between batches, where the distance matrix is based on Spearman correlation between median expression of highly variable features for the whole dataset, and the dendrogram was created via hierarchical clustering of batches on this correlation matrix.
**(L)** UMAP of all snRNA-seq nuclei colored by both target region and batch identity.
**(M)** Relative proportion of nuclei of each type for all snRNA-seq batches.
Brain diagrams were reproduced from Paxinos and Franklin (2005).[36]

#### Figure S7

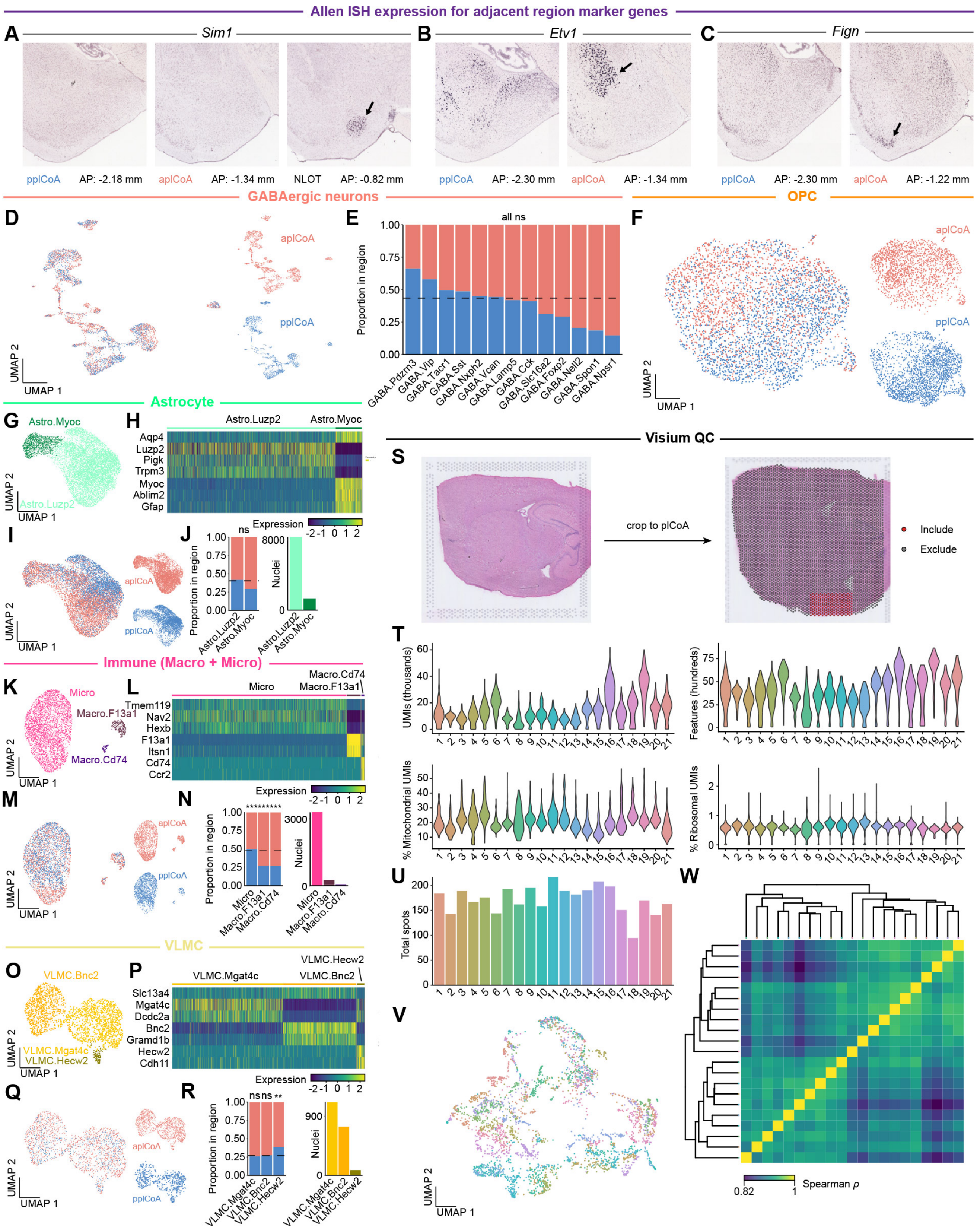

Allen ISH expression for adjacent region marker genes
A Sim1
B Etv1
C Fign
pplCoA AP: -2.18 mm
aplCoA AP: -1.34 mm
NLOT AP: -0.82 mm
pplCoA AP: -2.30 mm
aplCoA AP: -1.34 mm
pplCoA AP: -2.30 mm
aplCoA AP: -1.22 mm
GABAergic neurons
OPC
D
E
F
all ns
Proportion in region
GABA.Pdzrn3
GABA.Vip
GABA.Tacr1
GABA.Sst
GABA.Nxph2
GABA.Vcan
GABA.Lamp5
GABA.Cck
GABA.Slc16a2
GABA.Foxp2
GABA.Nell2
GABA.Spon1
GABA.Npsr1
UMAP 1
UMAP 2
Astrocyte
G
H
Astro.Myoc
Astro.Luzp2
Aqp4
Luzp2
Pigk
Trpm3
Myoc
Ablim2
Gfap
Expression
I
J
ns
Nuclei
Immune (Macro + Micro)
K
L
Micro
Macro.F13a1
Macro.Cd74
Tmem119
Nav2
Hexb
F13a1
Itsn1
Cd74
Ccr2
M
N
VLMC
O
P
VLMC.Bnc2
VLMC.Mgat4c
VLMC.Hecw2
Slc13a4
Mgat4c
Dcdc2a
Bnc2
Gramd1b
Hecw2
Cdh11
Q
R
ns ns **
Visium QC
S
crop to plCoA
Include
Exclude
T
UMIs (thousands)
Features (hundreds)
% Mitochondrial UMIs
% Ribosomal UMIs
U
Total spots
V
W
Spearman ρ
0.82
1

**Figure S7. Additional information and quality control for spatial gene expression.** Related to **Figure 4**.
(**A-C**) Allen ISH data for marker genes from molecular cell types adjacent to, but not within plCoA. *Sim1* (**A**) marks cells in the NLOT, *Etv1* (**B**) marks cells in the BLA and posterior basomedial amygdala, and *Fign* (**C**) marks cells in the cortex-amygdalar transition area, but none of these mark cells in the plCoA. Arrow points to respective clusters of cells marked with these genes.
(**D**) UMAP of all plCoA GABAergic neurons, colored by domain of origin.
(**E**) Relative proportion of molecular subtype nuclei from each domain within GABAergic neurons. Dotted line indicates chance level for plCoA GABAergic neuron nuclei.
(**F**) UMAP of all plCoA OPCs, colored by domain of origin.
(**G**) Two-dimensional UMAP of astrocytes, colored by molecular cell type.
(**H**) Heatmap of astrocyte subtype marker genes.
(**I**) UMAP of all plCoA astrocytes, colored by domain of origin.
(**J**) Left, relative proportion of molecular subtype nuclei from each domain within astrocytes. Dotted line indicates chance level for plCoA astrocyte nuclei. Right, relative abundance of each astrocyte subtype within plCoA.
(**K**) Two-dimensional UMAP of immune cells, colored by molecular cell type.
(**L**) Heatmap of immune cell subtype marker genes.
(**M**) UMAP of all plCoA immune cells, colored by domain of origin.
(**N**) Left, relative proportion of molecular subtype nuclei from each domain within immune cells. Dotted line indicates chance level for plCoA immune cell nuclei. Right, relative abundance of each immune cell type within plCoA.
(**O**) Two-dimensional UMAP of VLMC nuclei, colored by molecular cell type.
(**P**) Heatmap of VLMC subtype marker genes.
(**Q**) UMAP of all plCoA VLMC nuclei, colored by domain of origin.
(**R**) Left, relative proportion of molecular subtype nuclei from each domain within VLMCs. Dotted line indicates chance level for plCoA VLMC nuclei. Right, relative abundance of each VLMC subtype within plCoA.
(**S**) Left, representative section on a Visium slide capture area stained with hematoxylin and eosin. Right, representative section with capture spots overlaid (grey) and plCoA-overlapping spots highlighted (red).
(**T**) Violin plots of quality metrics for individual Visium sections on a per-spot basis in plCoA-overlapping spots ($N$ = 21 sections). Upper left, UMIs per spot; upper right, features per spot; lower left, proportion mitochondrial UMIs per spot; lower right, proportion ribosomal UMIs per spot.
(**U**) Number of plCoA-overlapping capture spots per section ($n$ = 3,616 total spots).
(**V**) Two-dimensional UMAP of all plCoA-overlapping capture spots, colored by section of origin.
(**W**) Evaluation of transcriptomic homology between sections, where the distance matrix is based on Spearman correlation between median expression of highly variable features for the whole dataset, and the dendrogram was created via hierarchical clustering of sections on this correlation matrix.
Across panels: ** $p < 0.01$; *** $p < 0.001$; ns, not significant. Additional specific details of statistical tests can be found in Supplemental Table 1.

#### Figure S8

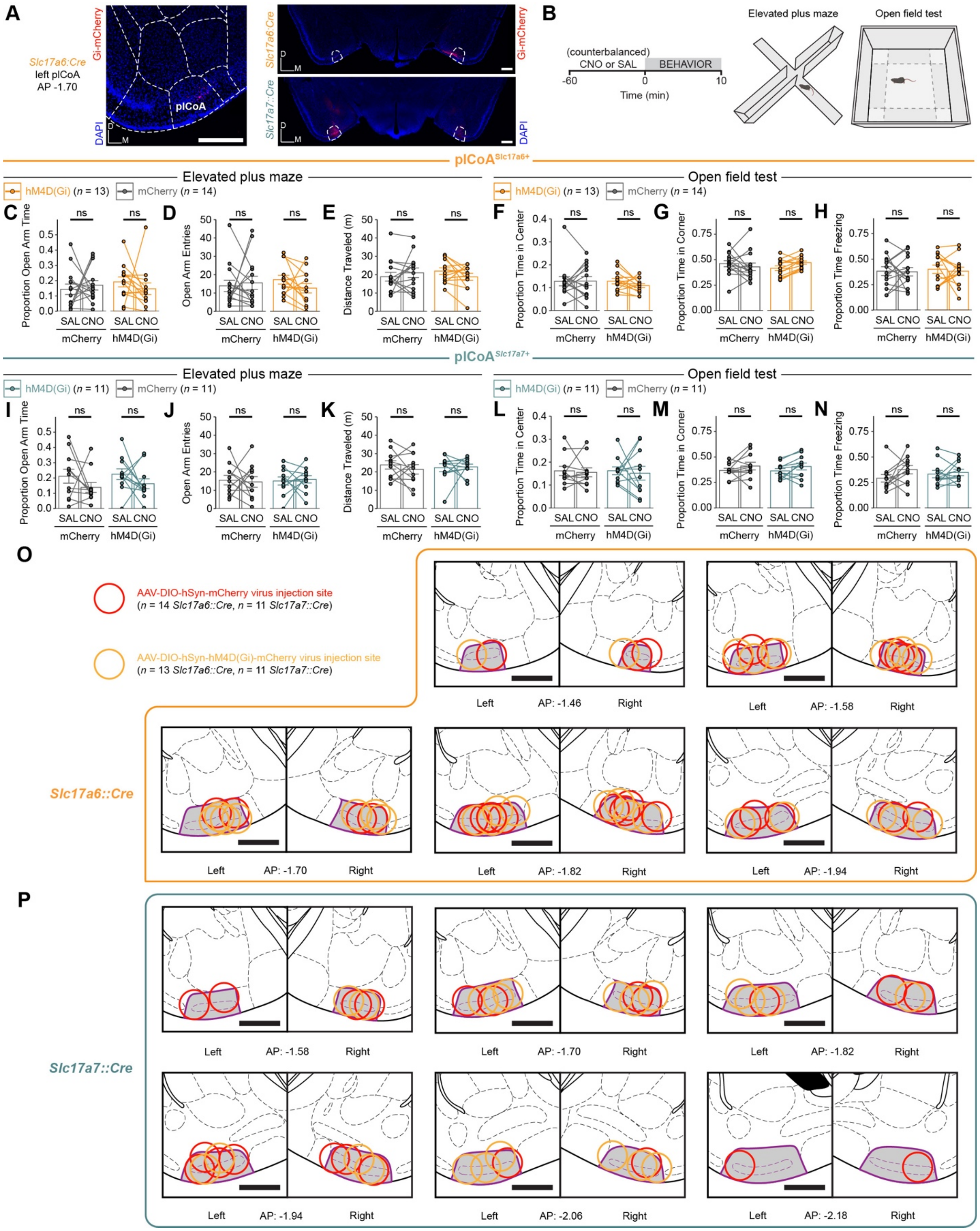
A
B
Slc17a6:Cre
left plCoA
AP -1.70
Gi-mCherry
DAPI
plCoA
Slc17a7::Cre
(counterbalanced)
CNO or SAL
BEHAVIOR
Time (min)
Elevated plus maze
Open field test
plCoA Slc17a6+
plCoA Slc17a7+
hM4D(Gi) (n = 13)
mCherry (n = 14)
hM4D(Gi) (n = 11)
mCherry (n = 11)
Proportion Open Arm Time
Open Arm Entries
Distance Traveled (m)
Proportion Time in Center
Proportion Time in Corner
Proportion Time Freezing
SAL CNO
mCherry
hM4D(Gi)
ns
O
P
AAV-DIO-hSyn-mCherry virus injection site
(n = 14 Slc17a6::Cre, n = 11 Slc17a7::Cre)
AAV-DIO-hSyn-hM4D(Gi)-mCherry virus injection site
(n = 13 Slc17a6::Cre, n = 11 Slc17a7::Cre)
Slc17a6::Cre
Slc17a7::Cre
Left
Right
AP: -1.46
AP: -1.58
AP: -1.70
AP: -1.82
AP: -1.94
AP: -2.06
AP: -2.18

**Figure S8. Additional information for Cre-dependent molecularly targeted chemogenetic inhibition experiments.** Related to **Figure 5**.
**(A)** Representative histology for inhibition experiments for AAV-hM4D(Gi)-mCherry-infected *Slc17a6::Cre* (left) or *Slc17a7::Cre* (right) animals.
(**B**) Strategy for chemogenetic inhibition in the open field test and elevated plus maze.
**(C-E)** Effect of chemogenetic inhibition of plCoA$^{Slc17a6+}$ neurons in the elevated plus maze. Inhibition does not induce a significant change in time spent (**C**) or number of entries (**D**) into the open arms, as well as distance traveled (**E**) in the elevated plus maze.
**(F-H)** Effect of chemogenetic inhibition of plCoA$^{Slc17a6+}$ neurons in the open field test. Inhibition does not induce a significant change in time spent in the center (**F**) or corners (**G**), as well as distance traveled (**H**) in the open field test.
**(I-K)** Effect of chemogenetic inhibition of plCoA$^{Slc17a7+}$ neurons in the elevated plus maze. Inhibition does not induce a significant change in time spent (**I**) and number of entries (**J**) into the open arms, as well as distance traveled (**K**) in the elevated plus maze.
**(L-N)** Effect of chemogenetic inhibition of plCoA$^{Slc17a7+}$ neurons in the open field test. Inhibition does not induce a significant change in time spent in the center (**L**) or corners (**M**), as well as distance traveled (**N**) in the open field test.
(**O**) Histologically verified placements for viral injection sites in *Slc17a6::Cre* animals infected with AAV-DIO-hSyn-mCherry (red) or AAV-DIO-hSyn-hM4D(Gi) (light orange) in molecularly targeted chemogenetic inhibition experiments.
(**P**) Same as (**O**), but in *Slc17a7::Cre* animals.
*n* denotes number of mice per group batched across 4-quad, elevated plus maze, and open field test experiments, exceeding *n* values for individual experiments due to behavioral cohort design (see STAR Methods). All mouse brain sections reproduced from Paxinos and Franklin, 5$^{th}$ Edition, with plCoA highlighted in grey and outlined in purple, and numbers below all images denote its anterior-posterior distance from bregma in this atlas.[36] All scale bars, 500 µm. Across panels: ns, not significant. Additional specific details of statistical tests can be found in Supplemental Table 1.

#### Figure S9

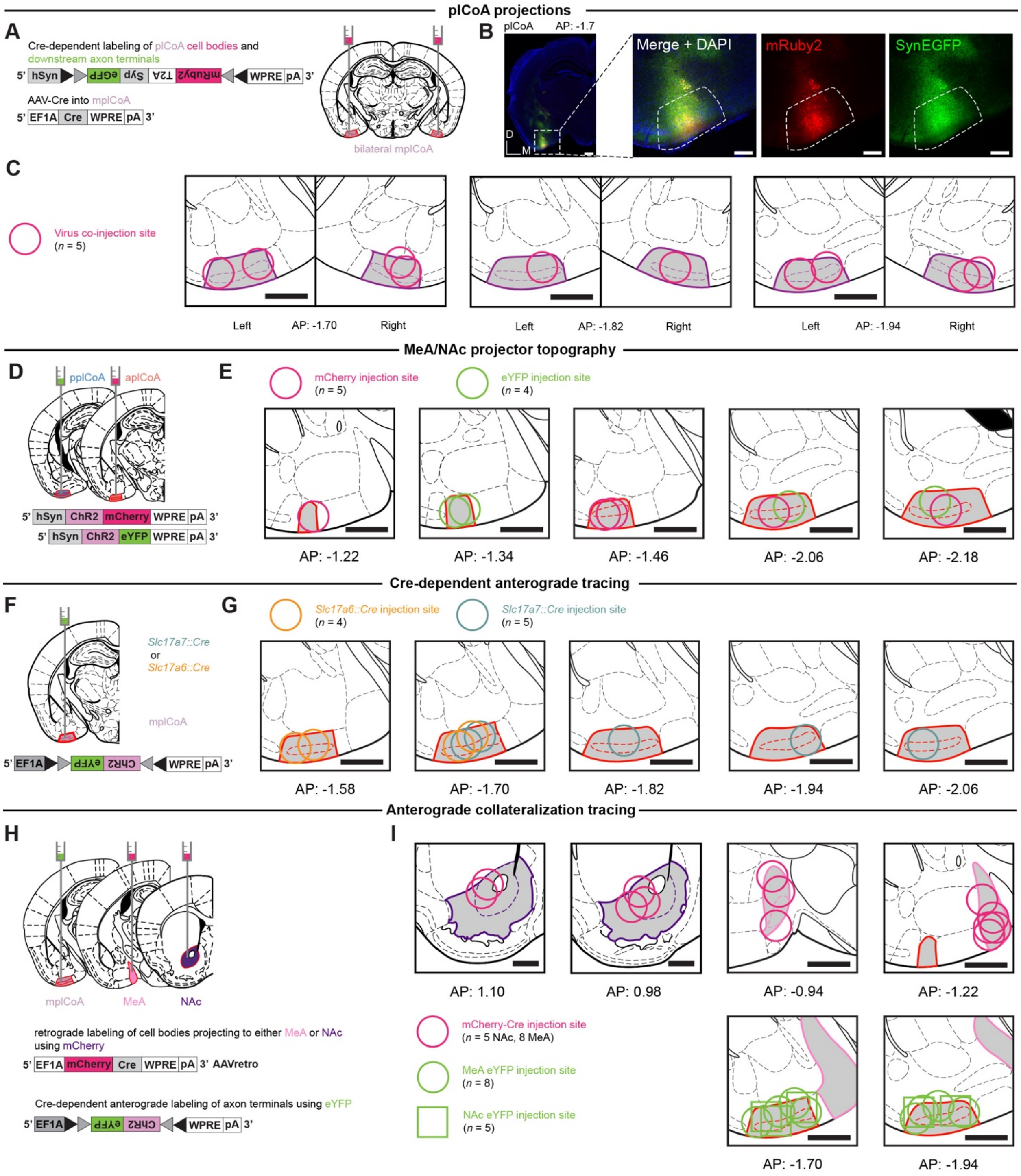
plCoA projections
A
Cre-dependent labeling of plCoA cell bodies and downstream axon terminals
5' hSyn eGFP Syp T2A mRuby2 WPRE pA 3'
AAV-Cre into mplCoA
5' EF1A Cre WPRE pA 3'
bilateral mplCoA
B
plCoA AP: -1.7
Merge + DAPI
mRuby2
SynEGFP
D
M
C
Virus co-injection site
(n = 5)
Left AP: -1.70 Right
Left AP: -1.82 Right
Left AP: -1.94 Right
MeA/NAc projector topography
D
pplCoA
aplCoA
5' hSyn ChR2 mCherry WPRE pA 3'
5' hSyn ChR2 eYFP WPRE pA 3'
E
mCherry injection site
(n = 5)
eYFP injection site
(n = 4)
AP: -1.22
AP: -1.34
AP: -1.46
AP: -2.06
AP: -2.18
Cre-dependent anterograde tracing
F
Slc17a7::Cre
or
Slc17a6::Cre
mplCoA
5' EF1A eYFP ChR2 WPRE pA 3'
G
Slc17a6::Cre injection site
(n = 4)
Slc17a7::Cre injection site
(n = 5)
AP: -1.58
AP: -1.70
AP: -1.82
AP: -1.94
AP: -2.06
Anterograde collateralization tracing
H
mplCoA
MeA
NAc
retrograde labeling of cell bodies projecting to either MeA or NAc using mCherry
5' EF1A mCherry Cre WPRE pA 3' AAVretro
Cre-dependent anterograde labeling of axon terminals using eYFP
5' EF1A eYFP ChR2 WPRE pA 3'
I
AP: 1.10
AP: 0.98
AP: -0.94
AP: -1.22
mCherry-Cre injection site
(n = 5 NAc, 8 MeA)
MeA eYFP injection site
(n = 8)
NAc eYFP injection site
(n = 5)
AP: -1.70
AP: -1.94

**Figure S9. Additional information for plCoA anterograde tracing experiments.** Related to **Figure 6**.

**(A)** Schematic for general anterograde output mapping strategy, where a virus labeling cell bodies in plCoA with mRuby and presynaptic terminals in downstream regions with synaptophysin-bound eYFP.

**(B)** Histological image of plCoA injection site in a representative animal. Scale bar, 200 μm.

**(C)** Histologically verified placements for viral injection sites in initial anterograde tracing experiments.

**(D)** Schematic for dual-color topographic anterograde tracing: two counterbalanced fluorophores were injected into aplCoA and pplCoA, and each color was quantified in major projection targets of plCoA.

**(E)** Histologically verified placements for viral injection sites in dual-color anterograde tracing experiments.

**(F)** Schematic for Cre-dependent anterograde output mapping strategy: a Cre-dependent virus expressing eYFP was injected into either an *Slc17a6::Cre* or *Slc17a7::Cre* animal to determine relative output enrichment for either broad cell type.

**(G)** Histologically verified placements for viral injection sites in Cre-dependent molecular anterograde tracing experiments.

**(H)** Schematic of anterograde viral strategy to explore collateralization of plCoA MeA and NAc projection neurons to other regions.

**(I)** Histologically verified placements for viral injection sites in collateralization experiments. Relevant regions are highlighted in grey and outlined: plCoA (red), NAc (purple, only in **I**), and MeA (pink, only in **I**). All mouse brain sections reproduced from Paxinos and Franklin, 5th Edition, and numbers below all images denote its anterior-posterior distance from bregma in this atlas.[36] All scale bars 500 μm unless noted elsewhere.

**Figure S10**

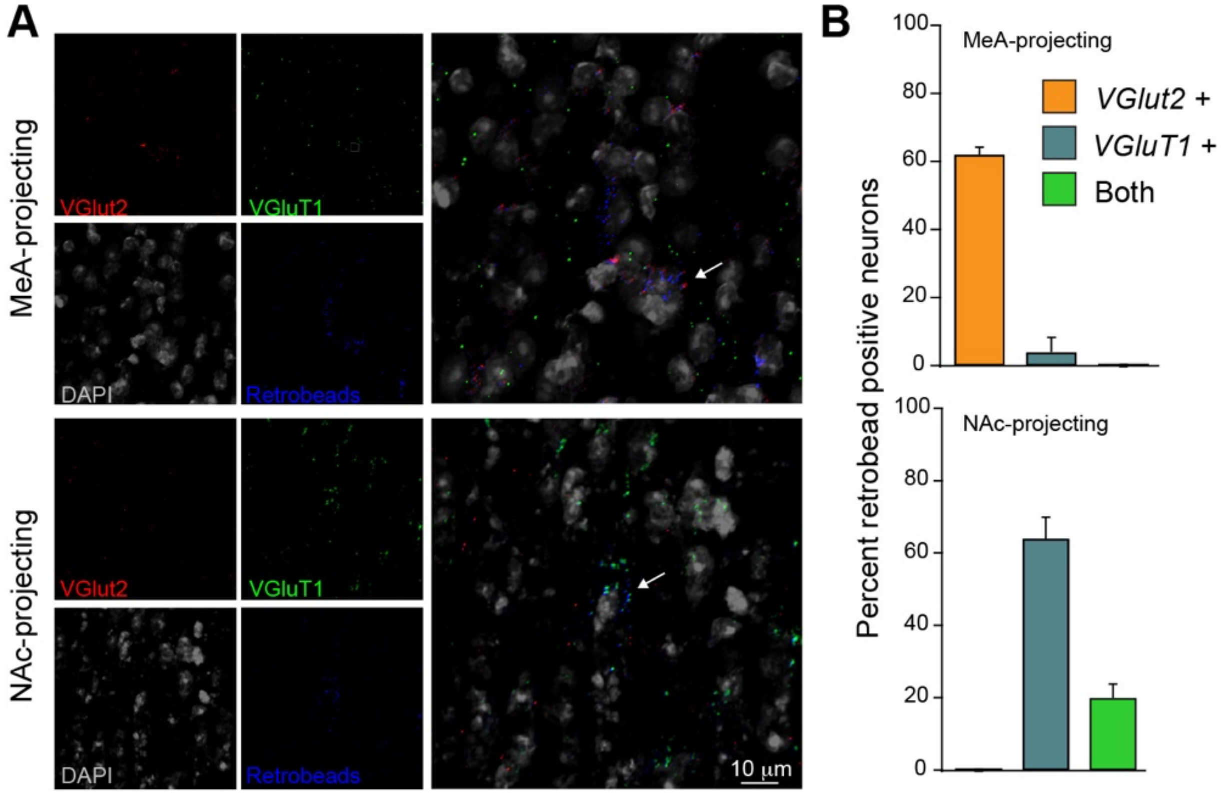


**Figure S10. Analysis of *VGluT* experession in MeA- and NAc-projecting neurons.** Related to Figure 6.

(**A**) Representative mplCoA images of *in situ* RNAscope labeling of *VGluT2* (red) and *VGluT1* (green) in animals with Retrobeads (blue) injected into the MeA (top) or NAc (bottom). The RNAscope and Retrobead signals are overlaid on DAPI staining (gray). White arrow in merged image points to a Retrobead+ and VGluT2+ cell.

(**B**) Quantification of the percentage of Retrobead labeled neurons positive for *VGluT1*, *VGluT2* or both for MeA- and NAc projecting neurons (top and bottom, respectively) in mplCoA. White arrow in merged image points to a Retrobead+ and VGluT1+ cell.

#### Figure S11

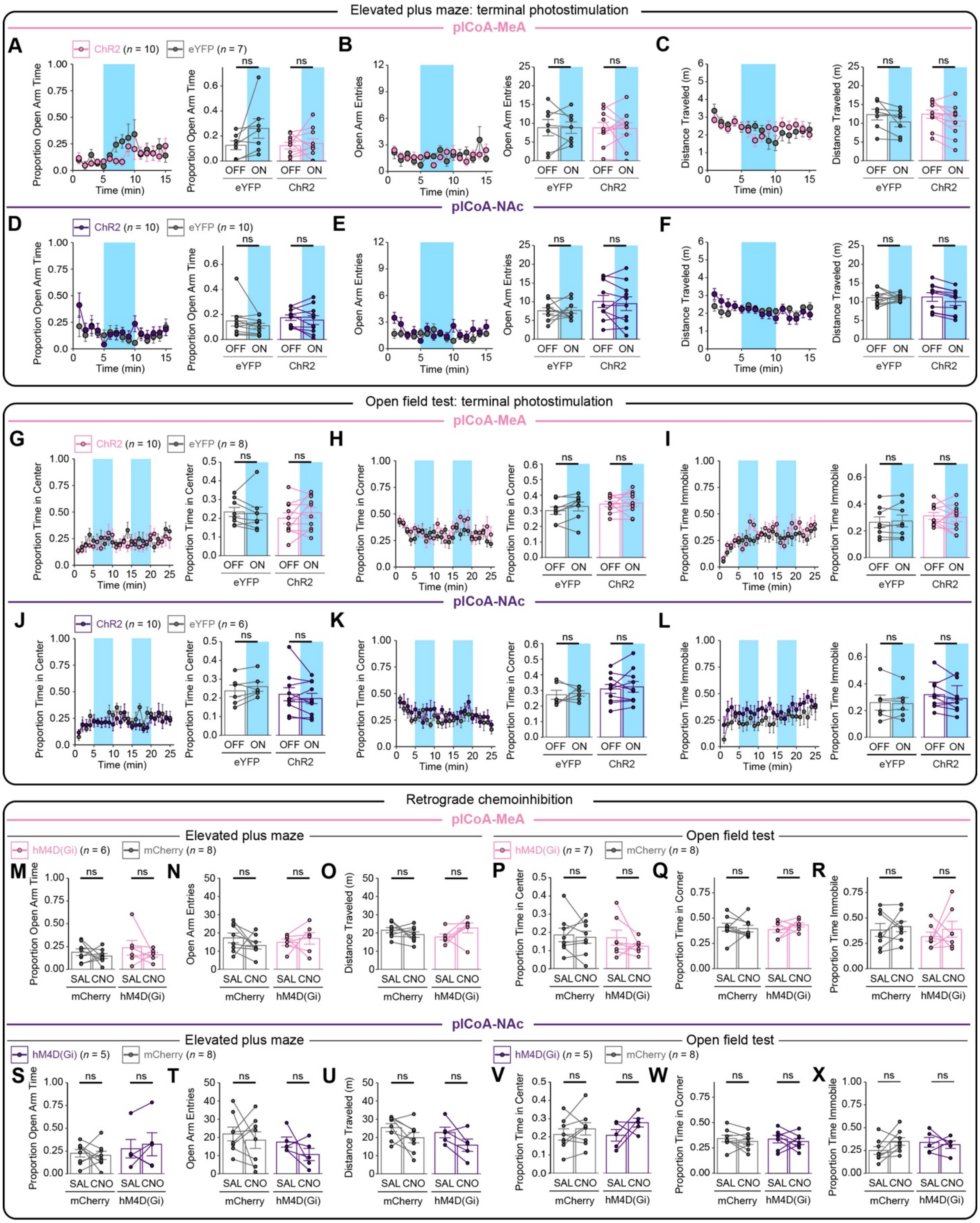
Elevated plus maze: terminal photostimulation
plCoA-MeA
A
ChR2 (n = 10) eYFP (n = 7)
Proportion Open Arm Time
Time (min)
ns
OFF ON
eYFP
ChR2
B
Open Arm Entries
C
Distance Traveled (m)
plCoA-NAc
D
ChR2 (n = 10) eYFP (n = 10)
E
F
Open field test: terminal photostimulation
G
ChR2 (n = 10) eYFP (n = 8)
Proportion Time in Center
H
Proportion Time in Corner
I
Proportion Time Immobile
J
ChR2 (n = 10) eYFP (n = 6)
K
L
Retrograde chemoinhibition
Elevated plus maze
hM4D(Gi) (n = 6) mCherry (n = 8)
Open field test
hM4D(Gi) (n = 7) mCherry (n = 8)
M N O P Q R
SAL CNO
mCherry
hM4D(Gi)
hM4D(Gi) (n = 5) mCherry (n = 8)
S T U V W X

**Figure S11. Manipulation of plCoA projections to MeA or NAc in either direction does not change features of behavior unrelated to innate valence.** Related to **Figure 7**.
**(A-C)** Effects of optogenetic stimulation of plCoA terminals in MeA in the elevated plus maze in time bins of 1 minute (left) and during off and on periods (right). Photostimulation does not induce a significant change in time spent (**A**) and number of entries (**B**) into the open arms, as well as distance traveled (**C**) in the elevated plus maze.
**(D-F)** Effects of optogenetic stimulation of plCoA terminals in NAc in the elevated plus maze in time bins of 1 minute (left) and during off and on periods (right). Photostimulation does not induce a significant change in time spent (**H**) and number of entries (**I**) into the open arms, as well as distance traveled (**J**) in the elevated plus maze.
**(G-I)** Effects of optogenetic stimulation of plCoA terminals in MeA in the open field test in time bins of 1 minute (left) and during off and on periods (right). Photostimulation does not induce a significant change time spent in the center (**G**) and corners (**H**), as well as distance traveled (**I**) in the open field test.
**(J-L)** Effects of optogenetic stimulation of plCoA terminals in NAc in the open field test in time bins of 1 minute (left) and during off and on periods (right). Photostimulation does not induce a significant change time spent in the center (**G**) and corners (**H**), as well as distance traveled (**I**) in the open field test.
(**M-O**) Effects of chemogenetic inhibition of plCoA-MeA projection neurons in the elevated plus maze. Inhibition does not induce a significant change in time spent (**M**) and number of entries (**N**) into the open arms, as well as distance traveled (**O**) in the elevated plus maze.
(**P-R**) Effects of chemogenetic inhibition of plCoA-MeA projection neurons in the open field test. Inhibition does not induce a significant change in time spent in the center (**P**) and corners (**Q**), as well as distance traveled (**R**) in the open field test.
(**S-U**) Effects of chemogenetic inhibition of plCoA-NAc projection neurons in the elevated plus maze. Inhibition does not induce a significant change in time spent (**S**) and number of entries (**T**) into the open arms, as well as distance traveled (**U**) in the elevated plus maze.
(**V-X**) Effects of chemogenetic inhibition of plCoA-NAc projection neurons in the open field test. Inhibition does not induce a significant change in time spent in the center (**V**) and corners (**W**), as well as distance traveled (**X**) in the open field test.
Across panels: ns, not significant. Additional specific details of statistical tests can be found in Supplemental Table 1.
